# Multi-omics resources for the Australian southern stuttering frog (*Mixophyes australis*) reveal assorted antimicrobial peptides

**DOI:** 10.1101/2023.09.22.558932

**Authors:** Simon Tang, Emma Peel, Katherine Belov, Carolyn J. Hogg, Katherine A. Farquharson

## Abstract

The number of genome-level resources for non-model species continues to rapidly expand. However, frog species remain underrepresented, with up to 90% of frog genera having no genomic or transcriptomic data. Here, we assemble the first genomic and transcriptomic resources for the recently described southern stuttering frog (*Mixophyes australis*). The southern stuttering frog is ground-dwelling, inhabiting naturally vegetated riverbanks in south-eastern Australia. Using PacBio HiFi long-read sequencing and Hi-C scaffolding, we generated a high-quality genome assembly, with a scaffold N50 of 369.3 Mb and 95.1% of the genome contained in twelve scaffolds. Using this assembly, we identified the mitochondrial genome, and assembled six tissue-specific transcriptomes. We also bioinformatically characterised novel sequences of two families of antimicrobial peptides (AMPs) in the southern stuttering frog, the cathelicidins and β-defensins. While traditional peptidomic approaches to peptide discovery have typically identified one or two AMPs in a frog species from skin secretions, our bioinformatic approach discovered 12 cathelicidins and two β-defensins that were expressed in a range of tissues. We investigated the novelty of the peptides and found diverse predicted activities. Our bioinformatic approach highlights the benefits of multi-omics resources in peptide discovery and contributes valuable genomic resources in an under-represented taxon.

## INTRODUCTION

Since the completion of the Human Genome Project in 2003, there has been a proliferation of high quality genomic and transcriptomic resources for non-model species (Hotaling et al. 2021; Lewin et al. 2022; Lewin et al. 2018). The ability to generate these ‘multi-omics’ resources in diverse species has clear comparative (Horgan and Kenny 2011), conservation (Formenti et al. 2022; Paez et al. 2022), and clinical benefits (Wong et al. 2021). As of July 2023, over 5,800 non-model animal genomes and 4,600 non-model animal transcriptomes are available in the largest global genome repository, the National Center for Biotechnology Information (NCBI) (Sayers et al. 2022). Despite the acceleration of genomic resources, only 46 genera within the Order Anura (10% of all frog genera) have genomic or transcriptomic resources available on NCBI (AmphibiaWeb 2022; Sayers et al. 2022). In Australia, despite 92% of frogs being endemic, only 4 of the 229 known endemic frog species currently have ‘omics level resources available (AmphibiaWeb 2022; Farquharson et al. 2023; Lamichhaney et al. 2021; Li et al. 2020; Rhie et al. 2021). The global disparity in genetic resources across taxa has been driven in part by the diverse structures of amphibian genomes and associated difficulties with sequencing and assembly (Lamichhaney et al. 2021; Smith et al. 2019). Frogs have highly variable genome sizes, (Lamichhaney et al. 2021; Liedtke et al. 2018); variable proportions of repetitive elements (Li et al. 2020; Seidl et al. 2019); and can have ploidy variation (Novikova et al. 2020).

An emerging application of genomic resources is the bioinformatic discovery of novel antimicrobial peptides (AMPs). AMPs are small peptides found across all classes of life that form an important part of the innate immune response (Hanson et al. 2019; Huan et al. 2020; Wang et al. 2016). The comprehensive characterisation of AMPs is valuable in understanding how different species respond to pathogens in their environment and their pleiotropic immune effects have also been explored as potential therapeutics, with several currently undergoing clinical trials (Mercer et al. 2020; Ridyard and Overhage 2021). Frogs are known to express AMPs from the exocrine glands of their skin in response to their exposure to diverse, microbial communities in their amphibious, freshwater habitats (Ladram and Nicolas 2016; Varga et al. 2018; Wang et al. 2020). Over 2,500 novel AMPs have been characterised from only 167 frog species on the Database of Anuran Defence Peptides v1.6 (DADP) (Novković et al. 2012). To date, frog AMPs have typically been characterised with a peptidomic approach; the iterative purification of AMPs showing antimicrobial activity from frog skin secretions using high-performance liquid chromatography (HPLC) (Yang et al. 2022). However, this method may not capture the suite of AMPs expressed in other tissues (Hao et al. 2012), nor AMPs with immune functions other than antimicrobial activity (He et al. 2019). In response, there has been increased interest in ‘mining’ genomic and transcriptomic resources for the genes that encode for AMPs (Peel et al. 2016). These methods are primarily homology-driven, using known sequences of large, well-characterised AMP families like cathelicidins and β-defensins to search for homologs (Dalla Valle et al. 2012). A genomics-driven approach has been applied to discover novel AMPs from a range of non-model species (Peel et al. 2016; Pérez de la Lastra et al. 2021; Wang et al. 2021; Yoo et al. 2015). However, despite the evolutionary divergence of Australian frogs, none have yet been investigated for AMPs using a bioinformatic approach.

The southern stuttering frog (*Mixophyes australis*) (Mahony et al. 2023) is a small, ground-dwelling frog inhabiting naturally vegetated riverbanks in south-eastern Australia, and is part of the Myobatrachidae family of endemic Australian ground frogs (Barker et al. 1995; Cogger 2000). The northern (*Mixophyes balbus*) and southern (*Mixophyes australis*) species of stuttering frog were once considered a single species, but were recently split (Mahony et al. 2023). Although yet to be officially assessed, application of the International Union for the Conservation of Nature (IUCN) threat assessment methods for the southern stuttering frog warrants a listing of ‘Endangered’ for this newly defined species (Mahony et al. 2023). As a ground-dwelling frog, southern stuttering frogs spend most of their life cycle exposed to large bodies of water and decomposing detritus, which may carry pathogens such as chytrid fungus (*Batrachochytrium dendrobatidis*) and Ranaviruses (Mahony et al. 2023; Murray and Hose 2005). We hypothesise that the selective pressures of the southern stuttering frog’s environment make it a good candidate for the discovery of novel and diverse AMPs. Preliminary studies on other *Mixophyes* species demonstrated some resistance against chytrid fungus in skin secretions (Hollanders et al. 2023; Woodhams et al. 2007). While these secretions have not been sequenced, these studies suggest possible AMP activity in the *Mixophyes* genus.

In this study, we use a combination of PacBio HiFi long-read, short-read, and Hi-C sequencing data to generate the first genomic and transcriptomic resources for the southern stuttering frog. Using these high-quality resources, we use bioinformatics to characterise and analyse the AMP families cathelicidins and β-defensins. By generating the first ‘omics level resources for the *Mixophyes* genus, this study provides a unique opportunity to investigate an otherwise largely understudied, Australian taxon.

## RESULTS

### Multi-omics resources for the southern stuttering frog

Using a combination of PacBio HiFi long-read and Hi-C sequencing data, we generated a high-quality genome assembly for the southern stuttering frog (Figure 1A). The genome is 3.13 Gbp in length, has a read coverage of 26× and a scaffold N50 of 369 Mbp (Table 1). Repeat-masking the genome revealed a high proportion of repetitive elements (53.7%; Table S1), within the broad expected range for Anuran genomes (32.0-77.1%; (Sayers et al. 2022)). The GC content of the genome, 40.3%, was also within the range of expected Anuran GC values (26.9-44.5%; (Sayers et al. 2022)). From the assembly, we identified a scaffold representing the mitochondrial genome, which contained 36 genes: 21 tRNAs, 2 rRNAs and 13 protein-coding genes (Figure 1B).

**Figure 1.**
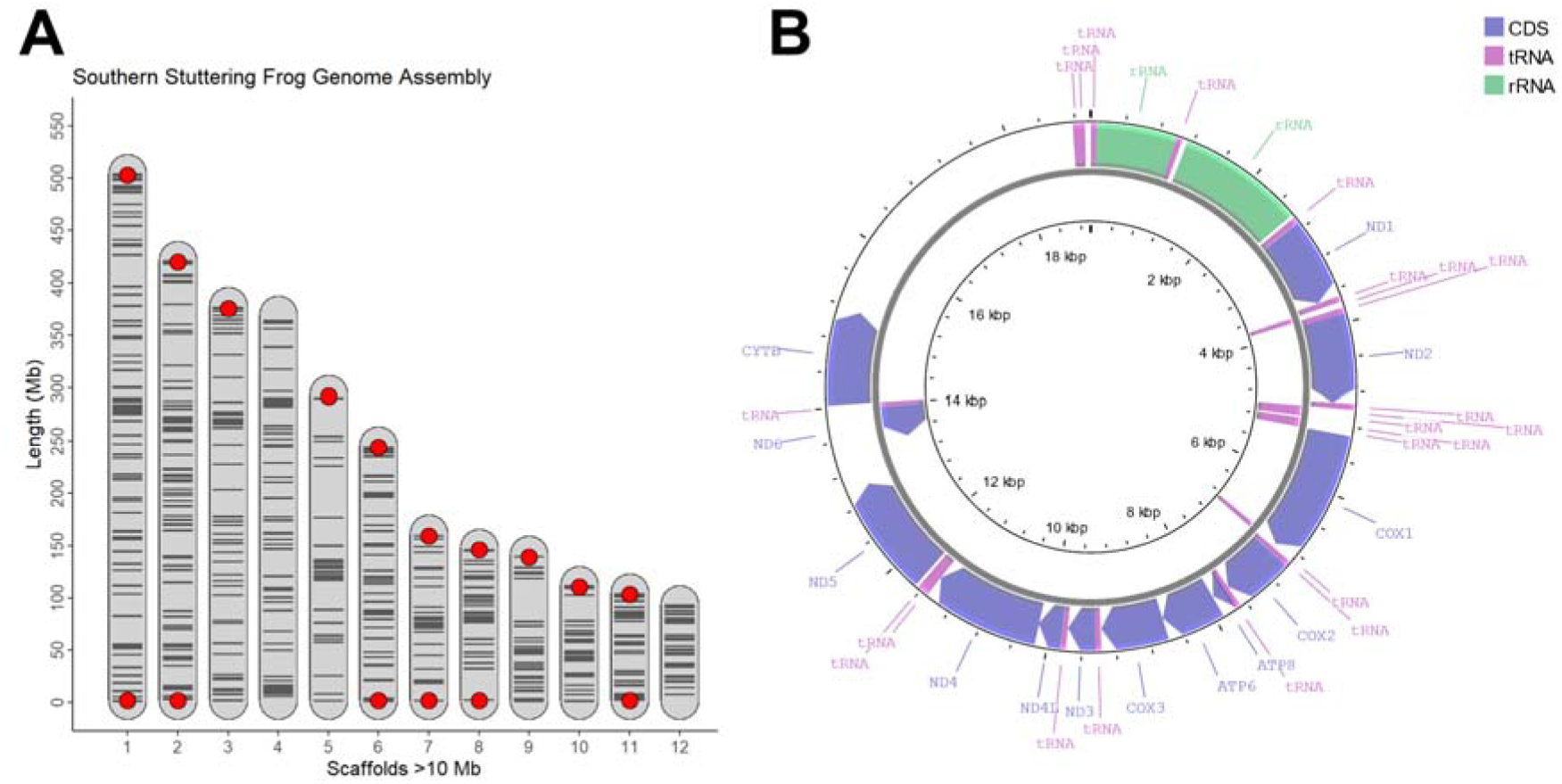
**A)** The 12 longest southern stuttering frog genome scaffolds, ordered by length. Red dots indicate the presence of telomeric sequences at the end of scaffolds. Horizontal, black lines indicate contig joins informed by the Hi-C sequencing data. **B)** Circos plot of the southern stuttering frog mitochondrial genome generated using Proksee (Grant et al. 2023). Outer ring contains genes on the forward strand, while the inner ring contains genes on the reverse strand.

**Table 1.** Statistics generated from the genome and transcriptomes of the southern stuttering frog (*Mixophyes australis*; this study), the model *Xenopus* species and three Australian frog genome assemblies. If available, the NCBI RefSeq or GenBank assembly number is provided. Read alignment rate refers to the percentage of tissue-specific RNA reads that mapped to the genome. BUSCO v5.3.2 (Simão et al. 2015) scores were calculated in this study for all genomes and transcriptomes using the vertebrata_odb10 lineage. There was no publicly available transcriptome for *L. dumerilii, P. ornatum* and *P. corroboree*, so transcriptome statistics could not be generated for these species. Full BUSCO scores for the genomes, transcriptomes and gene annotations are provided in Table S2.

| Genome Statistics |  |  |  |  |  |  |
| --- | --- | --- | --- | --- | --- | --- |
| Species and Assembly Number | Size (Gb) | Contig No. | Contig L/N50 (Mb) | Scaffold No. | Scaffold L/N50 (Mb) | Complete BUSCOs (%) |
| Southern stuttering frog (*Mixophyes australis*) | 3.120 | 1,290 | 85/11.04 | 790 | 4/369.31 | 91.3 |
| African clawed frog (*Xenopus laevis*) [GCF_017654675.1] | 2.742 | 716 | 35/22.45 | 55 | 8/155.25 | 95.9 |
| Western clawed frog (*Xenopus tropicalis*) [GCF_000004195.4] | 1.451 | 851 | 32/14.63 | 167 | 5/153.96 | 95.3 |
| Southern banjo frog (*Limnodynastes dumerilii*) [GCA_011038615.1] | 2.379 | 739,331 | 58,116/0.01 | 520,896 | 2,127/0.29 | 80.9 |
| Ornate burrowing frog (*Platyplectrum ornatum*) [GCA_016617825.1] | 1.065 | 238,568 | 39,409/0.004 | 148,035 | 9,283/0.03 | 42.4 |
| Corroboree frog (*Pseudophryne corroboree*) [GCA_028390025.1] | 8.873 | 5,826 | 353/6.82 | 3,127 | 5/846.89 | 93.1 |
| Transcriptome Statistics |  |  |  |  |  |  |
| Species | Type | Alignment Rate (%) | No. Transcripts | Complete BUSCOs (%) |  |  |
| Southern stuttering frog (*M. australis*) | Brain | 86.23 | 66,201 | 89.9 |  |  |
|  | Gonads (male) | 78.62 | 88,716 | 93.0 |  |  |
|  | Ventral skin | 88.32 | 52,141 | 89.3 |  |  |
|  | Dorsal skin | 87.96 | 55,849 | 89.8 |  |  |
|  | Liver | 89.37 | 46,579 | 84.0 |  |  |
|  | Spleen | 83.62 | 60,664 | 90.2 |  |  |
|  | Global | N/A | 192,115 | 96.1 |  |  |
| African clawed frog (*X. laevis*) [GJQV000000000.1] | Follicle | N/A | 333,966 | 70.8 |  |  |
| Western clawed frog (*X. tropicalis*) [GIVH000000000.1] | Gonads (male and female) | N/A | 372,630 | 95.7 |  |  |

Over 95% of the genome was assembled in twelve scaffolds (Figure 1A; Figure S1), which likely represent the chromosomes based on previous karyotypic analysis of other *Mixophyes* species (2n=24), including *M. hihihorlo* (Donnellan et al. 1990), *M. fasciolatus* and *M. schevilli* (Schmid et al. 2002). Six scaffolds were flanked at each end by long, telomere-like repeats, while four scaffolds had telomere-like repeats on a single end (Figure 1A). Additionally, a substantial drop in scaffold length between the 12^th^ and 13^th^ scaffold (94.13Mbp vs 4.75Mbp) was noted, further suggesting 12 chromosome-level scaffolds.

To compare the contiguity and completeness of the genome to other frog genomes, we calculated several genomic statistics, including benchmarking universal single-copy orthologs (BUSCO) (Simão et al. 2015) using the vertebrata_odb10 lineage. Genome statistics were calculated for two model frog species and the three publicly available scaffolded Australian frog genomes (Table 1). The southern stuttering frog had a similar scaffold L50 (L50 = 5) to other recent high-quality genome assemblies such as *Pseudophryne corroboree* (L50 = 4). Our BUSCO analysis revealed that our assembly contained 91.8% of complete BUSCOs, 2.9% were present but fragmented, while 5.3% were missing. The proportion of complete BUSCOs in the southern stuttering frog is slightly lower, but still comparable to the model reference *Xenopus laevis* (95.9%). An unbiased assessment of whole genome completeness with merqury (Rhie et al. 2020) showed that our genome assembly was highly complete (93.5%) and accurate (Q60+).

Using Illumina short-read RNA sequencing, we generated six reference-guided, tissue-specific transcriptome assemblies (dorsal skin, ventral skin, liver, spleen, brain, and gonads) that were aligned into a global transcriptome. All tissues had high mapping rates to the genome (78.62-89.37%), with both the tissue-specific and global transcriptomes yielding high complete BUSCO scores (84.0-96.1%; Table S2). A total of 36,540 genes were annotated using FGENESH++ (Solovyev et al. 2006). Of these, 16,355 were annotated using evidence from the coding regions of the global transcriptome, similar to the 16,279 predicted coding regions provided as input. A further 12,892 genes were annotated via homology to non-redundant, metazoan, proteins from NCBI, and 7,293 were annotated *ab initio*. Proteins from these annotated genes yielded a complete BUSCO score of 86.0%, comparable to the genome annotation of *Limnodynastes dumerilii*, and lower than the *Xenopus* spp. annotations (Table S2). However, a large proportion of the stuttering frog annotated genes were fragmented (8.5%), similar to *L. dumerilii* (8.0%).

### Characterisation of cathelicidins and ***β***-defensins

We used the genome and transcriptome assemblies of the southern stuttering frog to characterise two families of AMPs: cathelicidins and β-defensins. These families were selected due to their conservation across vertebrate species (Kosciuczuk et al. 2012; Tu et al. 2015), highly conserved structures and motifs (Chen et al. 2021; Shen et al. 2016; Yu et al. 2022) and demonstrated antimicrobial potency (Shen et al. 2016; Wei et al. 2015). We identified 12 cathelicidin (MA-CATH1-12) and two β-defensin (MA-BD1 and MA-BD2) genes in the genome using homology-based search strategies. Full-length transcripts of all cathelicidins and β-defensins were identified in the global transcriptome. All putative AMPs contained the characteristic features of each family, including expected exon number (four for cathelicidins and two for β-defensins), conserved amino acid residues, motifs, and domain structure (Figure S2 and S3). As expected, each AMP family was encoded in clusters within the genome; all cathelicidins were located on scaffold 5 and all β-defensins on scaffold 2 (Figure S4). Cathelicidin and β-defensin genes were named in the order in which they are encoded within the genome (Table S3; Figure S4). Amino acid sequences of all AMPs identified in this study, with predicted signal peptide and mature peptide domains are provided in Figures S2 and S3.

### Properties, evolutionary relationships and expression patterns of cathelicidins and ***β***-defensins

The mature peptide domains of cathelicidins and β-defensins are the bioactive, antimicrobial portion of the peptide. In the southern stuttering frog, the mature peptides of the cathelicidins ranged between 10-161 amino acids in length (Figure S2), while the two β-defensins were both 47 amino acids (Figure S3). The mature peptides of AMPs generally have a high positive charge which facilitates electrostatic interaction and attachment to microbial cell membranes (Huan et al. 2020). Stuttering frog cathelicidins MA-CATH1, 2 and 3 had a high cationic charge of 8.9, 4.9 and 27.3 respectively at pH 7 (Table S4). However, several AMPs identified in this study were weakly cationic (charge 0-3), and MA-CATH8 and MA-CATH2 were anionic. Similarly, while a large proportion (10/14) of the characterised AMPs had >30% hydrophobic residues, another general trend of AMPs (Chen and Lu 2020), some peptides like MA-CATH3 had as few as 12.42% hydrophobic residues (Table S4). Amphipathicity is often seen in AMPs to permeabilise microbial membranes (Chen and Lu 2020). Through visual inspection of Kyte and Dolittle hydropathicity plots, almost all the southern stuttering frog AMPs exhibited amphipathicity, with the N-terminus of the peptides being generally more hydrophilic than the C-terminus (Figure S5A, C). MA-CATH3 exhibited some regions of hydrophobicity, but was largely hydrophilic (Figure S5B).

As the first AMPs characterised in the *Mixophyes* genus, we explored their sequence diversity and relationship to other known frog AMPs. All stuttering frog AMPs displayed low percent identity to other known AMPs by BLAST, with MA-BD2 having the highest percent identity of all AMPs in this study to a Chinese spiny frog (*Quasipaa spinosa*) β-defensin (64.15%) (Table S4). As expected, maximum likelihood phylogenetic trees generated using all known frog cathelicidins and β-defensins reflected this diversity, as indicated by the long branch lengths (Figure 2). MA-CATH1-3 and MA-CATH4-8 formed two species-specific clades within the cathelicidin tree, some of which were strongly supported with >95% ultrafast bootstrap support (Figure 2). MA-CATH9-12 clustered within clades containing AMPs from multiple Asian and African frog species, albeit with low bootstrap support (Figure 2). While Figure 3 suggests that the southern stuttering frog β-defensins are more closely related than the other β-defensins, this relationship was not well supported.

**Figure 2.**
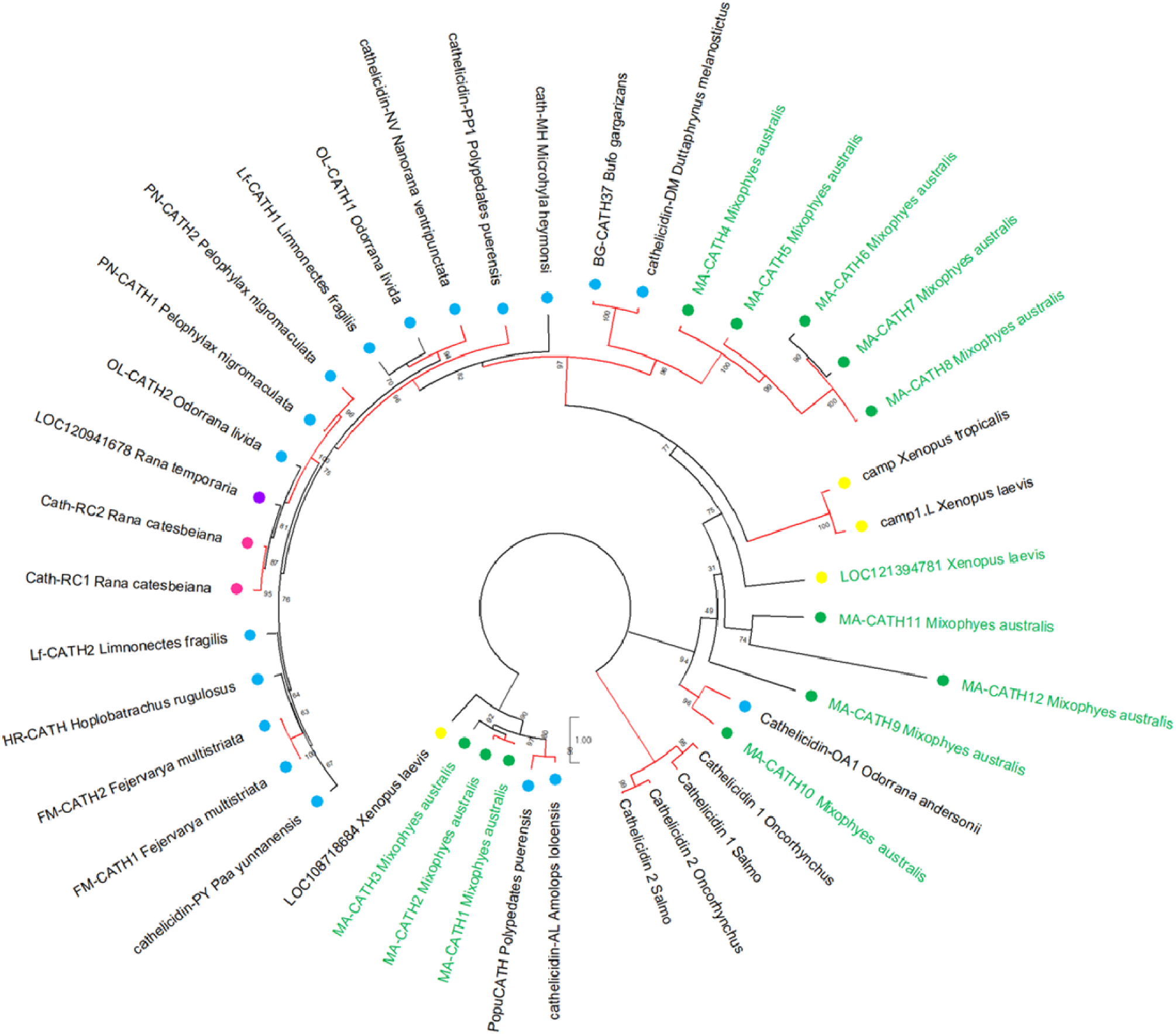
Phylogenetic relationships amongst frog cathelicidins. Tree was generated using the maximum likelihood method, and rooted using four fish cathelicidins as an outgroup. Branches are coloured by ultrafast bootstrap values, with values >95% in red, and the remaining branches coloured in black. Stuttering frog cathelicidins are coloured green. Cathelicidins are labelled by geographic region; Oceania is green; Asia is blue; North America is pink; Africa is yellow; and Asia and Europe combined is purple. The tree was annotated using MEGA11. For frog cathelicidin sequences used, see Table S5.

**Figure 3.**
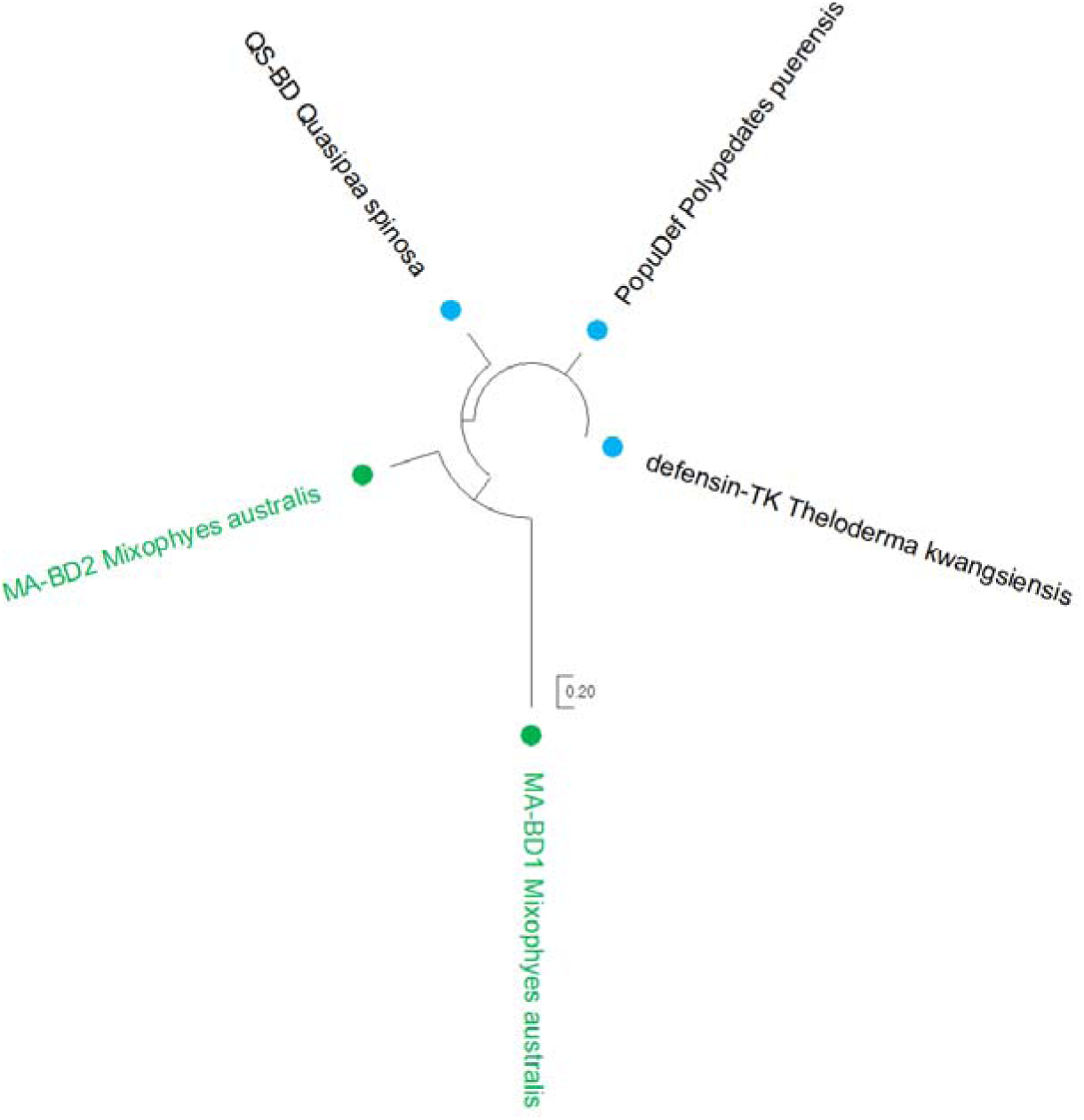
Phylogenetic relationships amongst frog defensins. Tree was generated using the maximum likelihood method and is unrooted. Branches are coloured by ultrafast bootstrap values, with values >95% in red, and the remaining branches coloured in black. Stuttering frog defensins are coloured green. Defensin are labelled by geographic region; Oceania is green and Asia is blue. The tree was annotated using MEGA11. For frog defensin sequences used, see Table S5.

Our comprehensive bioinformatic AMP discovery approach allowed us to explore the expression of AMPs across a range of tissues beyond skin, which is typically the target of peptidomic approaches to AMP discovery. As expected, we observed high AMP gene expression in the dorsal and ventral skin. However, we also observed AMP gene expression in other internal organs (Figure 4). AMP gene expression was found in the liver, spleen, and gonads, with MA-CATH6 and 8 showing the highest expression. MA-CATH10 and MA-BD2 exhibited higher expression in the skin than other AMPs, while some peptides like MA-CATH12 were lowly expressed across all the tissues when compared to other AMPs (Figure 4).

**Figure 4.**
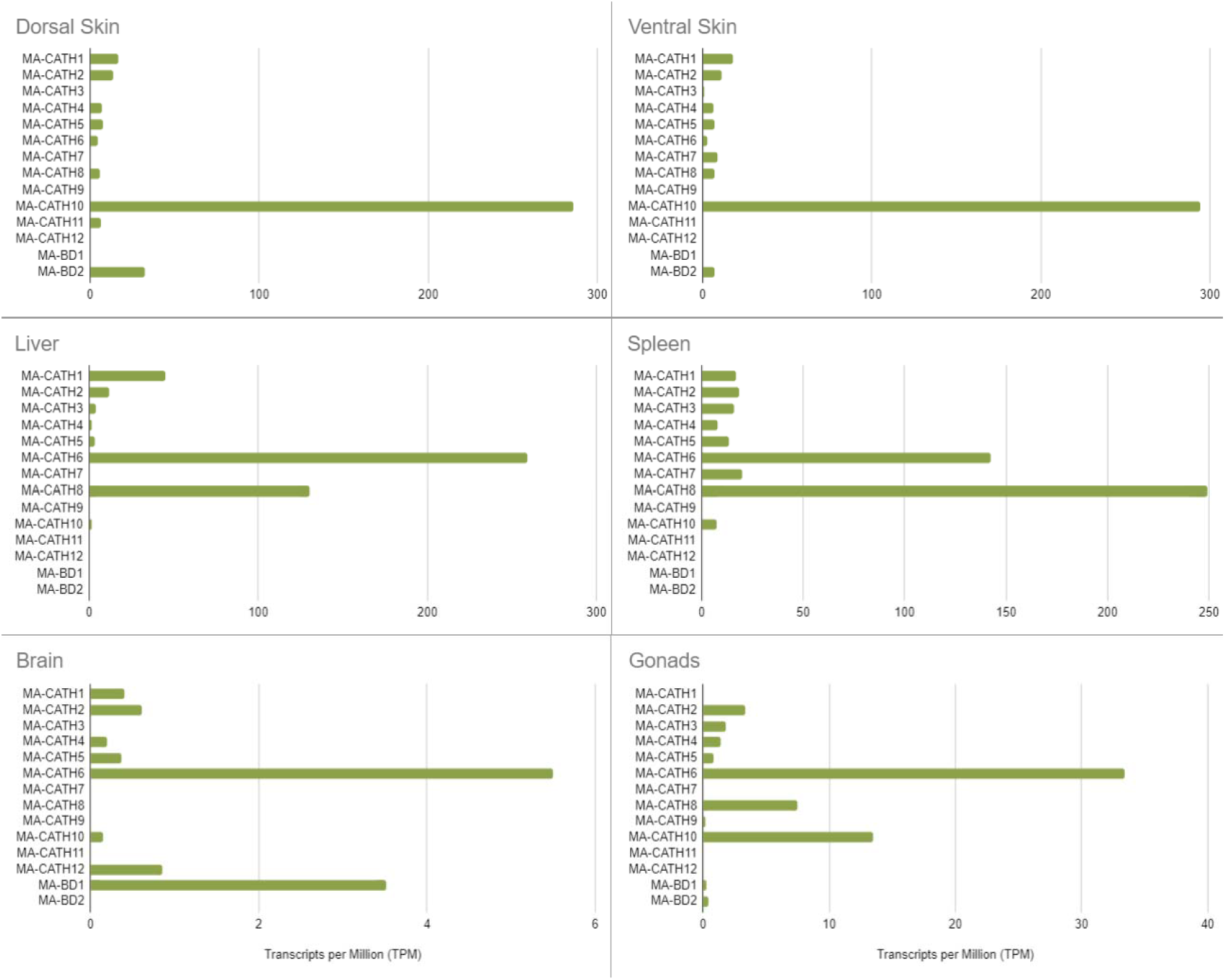
Expression data of the novel cathelicidins (MA-CATH1-12) and β-defensins (MA-BD1-2) from the stuttering frog. Transcripts per Million (TPM) values were derived from the tissue-specific transcriptomes. Note different x-axis scales due to differences in relative expression of AMPs between tissues.

## DISCUSSION

Here, we used a combination of PacBio HiFi and Hi-C data to generate the first contiguous, annotated, reference genome for the southern stuttering frog. In addition, we assembled the mitochondrial genome and six tissue-specific transcriptomes that were merged into a global transcriptome. The level of resources generated for this single species is comparable with model *Xenopus* species, as few frogs have a publicly available nuclear genome, mitochondrial genome, as well as transcriptome. Additionally, they are the first ‘omics level resources in the *Mixophyes* genus. As the Myobatrachidae family is one of the oldest, most diverse frog families in Australia (Brennan et al. 2023), these resources add to our understanding of Australian fauna.

To demonstrate the insights that can be gained from these genomic and transcriptomic resources, we bioinformatically characterised 12 cathelicidins and two β-defensins, the first Australian frog AMPs to be discovered using this approach. While other peptidomics-based studies describing cathelicidins in frogs have revealed at most two per species (Ling et al. 2014; Yu et al. 2013), 12 cathelicidins were discovered in this study. Similarly, while one β-defensin has been characterised per species in frogs (Shen et al. 2016; Yu et al. 2022), two were found in this study. The expression analysis from the transcriptomic data revealed some of these AMPs, such as MA-CATH1, MA-CATH10 and MA-BD2, were primarily expressed in the dorsal and ventral skin. However, several others had little to no expression in the skin but were primarily expressed in the liver (MA-CATH6), spleen (MA-CATH7-8), and low levels in the brain (MA-BD1). The peptides that were not expressed in the skin would not have been identified if the more conventional, peptidomics approach of screening skin secretions was used. Our results reveal the benefits of a genomic-based AMP discovery approach and the need for more amphibian genomic resources to characterise such peptides.

It is possible that the evolutionary isolation of Australian amphibians compared to other characterised species has resulted in the diverse suite of cathelicidins and β-defensins observed. Alternatively, the range of cathelicidins and β-defensins characterised here may suggest that the southern stuttering frog is under stronger microbial exposure than other frogs. A previous study on *Diptera* (fly) species has shown that AMP diversity and gene duplication is positively correlated with microbial exposure (Hanson et al. 2019). This is also likely the case in frogs as they adapt to different global aquatic and terrestrial environments. It is more likely, however, that other frog species and genera may also exhibit a similar number and diversity of cathelicidins and β-defensins which are uncharacterised. The lack of bioinformatic AMP discovery for most frog species makes comparative analysis difficult. Future investigations into frog AMP diversity that incorporate a genomic-based discovery platform will facilitate direct comparisons between species.

Our phylogenetic analysis revealed that some southern stuttering frog cathelicidins formed species-specific sister clades to those containing cathelicidins from frogs of Europe, North America, and Asia (Figure 2). This may indicate that these particular cathelicidins were the result of gene duplication events that occurred after the southern stuttering frogs diverged from the other frog species within the phylogenetic tree. Indeed, several southern stuttering frog cathelicidins have the same signal and cathelin region, but a different mature peptide. Other southern stuttering frog cathelicidins clustered with cathelicidins from European and African frog species (Figure 2). This suggests these cathelicidins may be the result of gene duplication events in a more distant common ancestor. However, due in part to the great variability in AMP sequences, many of the relationships identified were not strongly supported, particularly for β-defensins, limiting the validity of insights drawn from these trees. It also remains difficult to ascertain whether the species-specific clusters of southern stuttering frog cathelicidins are truly unique to *M. australis* or the *Mixophyes* genus more broadly, as there are currently no other characterised cathelicidins from *Mixophyes* frogs. Additionally, the number of known cathelicidins and β-defensins across the Anuran order is limited, with only one cathelicidin characterised in a European frog, and no known cathelicidins or β-defensins from South American frog species. Evolutionary relationships across these geographical regions are likely not captured. As more frog cathelicidins and β-defensins are characterised, future investigations may be able to better identify the evolutionary patterns of AMP diversity across all frogs.

The characterised and predicted properties of the stuttering frog AMPs suggest that they may play diverse immunological roles. AMPs are generally cationic and amphipathic due to their electrostatic interactions with anionic glycolipids on prokaryotic membranes, which facilitate membrane permeability and cell lysis (Hancock and Rozek 2002; Rivas et al. 2009). However, AMPs can also exhibit diverse activities beyond antimicrobial activity and may serve other immune functions. For example, an anionic cathelicidin from a salamander species (TK-CATH) had no tested antimicrobial activity, but instead inhibited pro-inflammatory cytokine gene expression when added to mammalian macrophage cell lines (Luo et al. 2021). Two of our AMPs were anionic (MA-CATH8 and MA-BD2; Table S4), suggesting that they may have other immune functions. Future investigations will need to validate these *in silico* findings, such as by synthesising these AMPs and investigating their effects on immune gene expression in a range of cell lines.

While our bioinformatic characterisation of novel AMPs has demonstrated one application of the newly generated genomic and transcriptomic resources, there are numerous other potential applications. For instance, custom DNA metabarcoding markers generated from the mitochondrial genome can now be developed for the southern stuttering frog, contributing to applied conservation outcomes in this threatened species. Metabarcoding has been extensively used to characterise the biodiversity of different environments (Ruppert et al. 2019). Mitochondrial metagenomics (mtMG) has been previously used to distinguish between closely related species of nematodes (Gendron et al. 2023). As the southern stuttering frog is a recently defined species, closely related to the northern population of stuttering frogs (*M. balbus*) (Mahony et al. 2023), the mitochondrial genome generated in this study may be a useful monitoring tool, particularly in defining their range and overlap if applicable. Highly contiguous genomes and transcriptomes can also be used to characterise genomic regions that have high repeat content, such as the major histocompatibility complex (MHC) (Vekemans et al. 2021). In frogs, the upregulation of MHC class I and II genes in the Montane brown frog (*Rana ornativentris*) has a functional role in tadpole development (Lau et al. 2020). MHC heterozygosity is also a significant predictor of chytrid fungus resistance in the *Lithobates* genus (Savage and Zamudio 2011; Trujillo et al. 2021). Finally, these resources may be used comparatively with other genomes to investigate conserved and specialised traits across taxa. For example, emerging consortia like the Zoonomia Project have made significant progress in advancing our understanding of mammalian adaptations and evolutionary history (Christmas et al. 2023). Similar efforts in Anurans would advance our understanding of a wealth of unique traits; some frog species can survive in extreme temperatures and environments (Larson et al. 2014; van Beurden 1980), produce a myriad of toxins (Raaymakers et al. 2017), and regenerate lost appendages as tadpoles (Aztekin et al. 2019). The incorporation of the southern stuttering frog genome in future studies into amphibians will facilitate a better representation of evolutionarily unique Australian biodiversity in these investigations. In short, generating high-quality multi-omics resources facilitates a plethora of investigations into the southern stuttering frog and amphibians at large.

## METHODS

### Sampling, Extractions and Sequencing

A wild caught, adult, male stuttering frog was medically euthanised in September 2021 (32°59’52.4’S 151°24’27.3’E). Heart and kidney tissue was flash frozen in liquid nitrogen for DNA extraction. Gonads, brain, dorsal skin, ventral skin, liver, and spleen tissue were stored in RNAlater at −80°C until RNA extraction. Lethal sampling was conducted under the University of Newcastle Animal Care and Ethics Committee (ACEC Number A-2013-339) and NSW scientific licence (SL190).

DNA was extracted from heart and kidney tissue using a Nanobind Tissue Big DNA Kit (Circulomics). Total extracted DNA was verified to be >20μg through Qubit fluorometric quantification (ThermoFisher Scientific). DNA was pooled and sequenced at the Australian Genome Research Facility (Brisbane, Queensland, Australia) using a SMRTbell® prep kit 3.0 (PacBio), and circular consensus sequencing (CCS) was performed using three SMRT cells on a PacBio Sequel II system. For Hi-C sequencing, heart and kidney tissue were washed twice for 5 minutes with 1×PBS using a rotator wheel at room temperature. Tissues were sequenced at the Biomolecular Resource Facility (Canberra, ACT, Australia), using the Arima Hi-C kit and sequenced as 150-bp paired-end (PE) reads on an Illumina NovaSeq 6000.

RNA was extracted from the six tissues using a Qiagen RNeasy Mini Kit. Concentrations of each sample were confirmed to be ≥25ng/µl using a Nanodrop spectrophotometer (ThermoFisher Scientific) and the RNA integrity number (RIN) measured using the standard Agilent RNA 6000 Nano Kit Protocol and BioAnalyzer (Agilent Technologies). Extractions <7 RIN were not sequenced. Extracted RNA was prepared at the Ramaciotti Centre for Genomics (Sydney, NSW, Australia), using the Illumina Stranded mRNA Prep Protocol and sequenced as 100-bp PE reads on an Illumina NovaSeq 6000 S1 flowcell.

### De novo Genome Assembly

Over 15 million raw HiFi reads were generated from three SMRT cells. To prevent low quality reads introducing errors to the assembly, reads with Phred (Q) quality score <20 were filtered out using bamtools v2.4.1 (Barnett et al. 2011). Reads containing adapter sequence were removed by HiFiAdapterFilt v2.0.0 (Sim et al. 2022) with default parameters. Further details on the computational requirements and estimated run times for all bioinformatic analyses are provided in Table S6. Hifiasm (Cheng et al. 2021) assembled the remaining reads, alongside paired Hi-C reads, into contigs. To merge the contigs into a scaffolded assembly, we first mapped the Hi-C reads to the unscaffolded genome following the Arima Hi-C mapping pipeline (A160156 v02; https://github.com/ArimaGenomics/mapping_pipeline). YaHS (Zhou et al. 2022) was used to merge contigs containing complementary pairs of reads, and the contact map visualised with Juicebox (Durand et al. 2016). One misassembly in the first scaffold was manually corrected. We used MitoHiFi v3.2 to identify and annotate the mitochondrial genome using a closely-related mitogenome as input (*Lechriodus melanopyga*; NC_019999.1) (Allio et al. 2020; Uliano-Silva et al. 2023).

Genome statistics (e.g., N50 and L50 values), were calculated with bbmap v38.86 (https://sourceforge.net/projects/bbmap/). We identified regions matching canonical telomere hexamer repeats (TTAGGG/CCCTAA) using FindTelomeres (https://github.com/JanaSperschneider/FindTelomeres). Benchmarking Universal Single-Copy Orthologs (BUSCO) v5.3.2 (Simão et al. 2015) analysis was performed on Galaxy Australia (Afgan et al. 2018), using the ‘genome’ mode, applying the ‘augustus’ gene-finding setting, and with the vertebrata_odb10 lineage. To compare the gene completeness of the stuttering frog with other genomes, BUSCO analysis was also performed with the same settings on two model frog genomes, the African clawed frog (*Xenopus laevis*; GCA_017654675.1) and the western clawed frog (*Xenopus tropicalis*; GCA_000004195.4), as well as the three publicly available scaffolded Australian frog genomes, the southern banjo frog (*Limnodynastes dumerilii*; GCA_011038615.1), ornate burrowing frog (*Platyplectrum ornatum*; GCA_016617825.1), and corroboree frog (*Pseudophryne corroboree*; GCA_028390025.1). An alternative assessment of genome completeness, inclusive of non-coding and repetitive regions, was performed for the stuttering frog and *P. ornatum* genomes using Merqury v1.3 (Rhie et al. 2020). RepeatModeler v2.0.1 (Flynn et al. 2020) was used to generate a *de novo* database of the repetitive regions in the stuttering frog genome. We characterised repeats and masked the genome with RepeatMasker v4.0.6 (Smit et al. 2013-2015). The repeat-masked genome was indexed with hisat2 v2.1.0 (Kim et al. 2015).

### Reference-aligned global transcriptome assembly

Over 900 million PE reads were generated across the six tissue transcriptomes. Low quality sequence calls, adapter and primer sequences were trimmed using Trimmomatic v0.39 (Bolger et al. 2014) with ILLUMINACLIP:TruSeq3-PE.fa:2:30:10, SLIDINGWINDOW:4:5, LEADING:5, TRAILING:5, and MINLEN:25 settings. The reads from each tissue sample were aligned to the genome with hisat2 v2.1.0 (Kim et al. 2015). StringTie v2.1.6 (Pertea et al. 2015) was used to merge aligned reads into tissue-specific transcriptomes. We used Transcriptome Annotation by Modular Algorithms (TAMA) v1.0 (Kuo et al. 2020) to combine the tissue-specific transcriptomes into a global transcriptome with adjustments to minimise duplicate transcripts. Briefly, the ‘-d merge_dup’ flag was applied to merge identical transcripts, and the ‘-z 500’ flag was applied to facilitate transcripts with variable 3’ ends (differences of up to 500bp) to be merged. Transcripts with weak evidence were removed, including transcripts that were found in only one tissue and were lowly expressed (fragments per kilobase of transcript per million fragments mapped [FPKM] < 0.1). CPC2 v2019-11-19 (Kang et al. 2017) was used to predict coding regions; non-coding regions were removed. Open reading frames were predicted by TransDecoder v2.0.1 (Haas et al. 2013). BUSCO v5.3.2 (Simão et al. 2015) analysis was performed using ‘transcriptome’ mode for both tissue-specific transcriptomes and the global transcriptome, as well as the *X. tropicalis* transcriptome for comparison.

### Genome Annotation

The genome was annotated with FGENESH++ v7.2.2 (Solovyev et al. 2006) informed by the global transcriptome. Non-mammalian settings were applied throughout the pipeline, and parameters were optimised for Anuran gene discovery by providing the *Xenopus* gene-finding matrix (Softberry). BUSCO v5.3.2 (Simão et al. 2015) analysis was performed on the annotation using ‘protein’ mode, as well as for the *X. laevis, X. tropicalis* and *L. dumerilii* annotations. The number of genes, exons and introns from the stuttering frog annotated assembly was calculated using the ‘genestats’ script (https://github.com/darencard/GenomeAnnotation/blob/master/genestats).

### AMP characterisation

To ensure a comprehensive search of genomic regions that could encode cathelicidins or β-defensins, the annotated genome and global transcriptome were queried using Basic Local Alignment Search Tool (BLAST) v2.2.30+ (Altschul et al. 1990; Altschul et al. 1997) and HMMER v3.3 (Eddy 2011). A dual program approach has previously been applied in other bioinformatic searches for cathelicidins and β-defensins (Helbing et al. 2019; Schutte et al. 2002; Whittington et al. 2008). Further explanation of the approaches used is provided in the Supplementary Extended Methods.

### AMP phylogeny

An unrooted phylogenetic tree was generated for the cathelicidin and β-defensin gene families. We first aligned amino acid sequences for each gene family through ClustalW alignments. Both alignments included the full prepropeptide sequences from the stuttering frog and other available frogs (Table S5). The cathelicidin alignment also included four fish cathelicidins as an outgroup (Table S5). The best-fitting substitution models for each alignment were selected using the ModelFinder option in IQ-TREE v2.2.0 (Minh et al. 2020) according to the Bayesian information criterion (Kalyaanamoorthy et al. 2017). For the cathelicidin alignment, the Jones Taylor Thornton (JTT) substitution model (Jones et al. 1992) was optimal, incorporating Invariant Sites (+I) and four components of gamma rate heterogeneity (+G4). For the β-defensin alignment, the Dayhoff model was optimal (Dayhoff et al. 1978). Maximum likelihood analysis was conducted in IQ-TREE v2.2.0 (Minh et al. 2020) with node support estimated by ultrafast bootstrap approximation with 1000 replicates (Hoang et al. 2017). The results were visualised and annotated using MEGA11 (Tamura et al. 2021).

### Characterisation and Prediction of AMP Structure, Properties and Expression

Cathelicidins are made up of a signal peptide, a conserved cathelin pro-region, as well as the mature, bioactive peptide (Chen et al. 2021; Tomasinsig and Zanetti 2005). To predict the signal peptide from the full prepropeptide sequence, the SignalP v6.0 webserver was used (Teufel et al. 2022) with the ‘Eukarya’ organism setting and the ‘Slow’ model mode. While the enzyme used to cleave the mature peptide from the cathelin pro-region is not known for amphibians, there is experimental evidence to support proprotein convertases or trypsin-like proteases that cleave at dibasic residues like lysine (K) and arginine (R) (Mu et al. 2017; Shi et al. 2020; Yu et al. 2013). Therefore, a two-tiered approach predicted the mature peptide of the novel cathelicidins. Proprotein convertase cleavage sites were first predicted from the last exon of the cathelicidins using ProP v1.0 (Duckert et al. 2004). If no cleavage was identified, trypsin cleavage sites were predicted using ExPASy’s peptide cutter tool (Gasteiger et al. 2003). For β-defensins, which generally consist of a signal peptide and a mature, bioactive peptide (Huttner and Bevins 1999; Semple and Dorin 2012; Tang and Selsted 1993), the signal peptide was predicted using SignalP, and the remaining peptide was annotated as the mature peptide.

The molecular weight and charge at pH 7 for each putative stuttering frog cathelicidin and β-defensin was calculated using Protein Calculator v3.4 (https://protcalc.sourceforge.net/). The percentage of hydrophobic residues in the AMPs was calculated on Peptide v2.0 (https://www.peptide2.com/) using the ‘Peptide Hydrophobicity/Hydrophilicity Analysis’ tool. The amphipathicity of the AMPs was determined by generating Kyte and Doolittle hydropathicity plots on ExPASy using Protscale (Gasteiger et al. 2003), with a window size of 5. These plots were inspected for the presence of ‘peaks’ and ‘troughs’, which indicate sections in the AMP of high and low hydropathicity (Peel et al. 2017). For the expression analysis, the Transcripts Per Million (TPM) values for each AMP was generated from the tissue-specific transcriptomes using StringTie v2.1.6 (Pertea et al. 2015).

## DATA ACCESS

The genome assembly, mitochondrial genome, and raw transcriptome sequencing data generated in this study have been submitted to the NCBI BioProject database (https://www.ncbi.nlm.nih.gov/bioproject/) under accession number PRJNA991157. Raw genome sequencing data is publicly available through the Bioplatforms Australia Threatened Species Initiative (https://data.bioplatforms.com/organization/threatened-species). The global transcriptome and annotation generated in this study are available on Amazon Web Services Australasian Genomes Open Data Store (https://awgg-lab.github.io/australasiangenomes/genomes.html).

## ACKNOWLEDGMENTS

We acknowledge the traditional custodians of the land where the specimen was collected, the Awabakal people, and pay respects to their elders past and present, and their ongoing connection to country. Thank you to M. Mahony, University of Newcastle, who collected the specimen. Thanks also to Elspeth McLennan for assistance with the DNA and RNA extractions. Computational resources were provided by Amazon Web Services and RONIN; the Australian FGENESH++ Service provided by the Australian Biocommons and the Pawsey Supercomputing Research Centre with funding from the Australian Government and the Government of Western Australia; Galaxy Australia, a service provided by the Australian Biocommons and its partners; and the University of Sydney’s High Performance Computing facility Artemis provided by the Sydney Informatics Hub. The authors wish to acknowledge the use of the services and facilities of the Australian Genome Research Facility; the Ramaciotti Centre for Genomics, UNSW; and the Biomolecular Resource Facility, Canberra. Funding was provided by the Threatened Species Initiative through a grant from the Australian Government Bushfire Recovery Scheme (ERF-WRR2-020), and the Australian Research Council Centre of Excellence for Innovations in Peptide and Protein Science (CE200100012).

## AUTHOR CONTRIBUTIONS

The project was conceived, and funding sought, by CJH and KB. ST undertook the genome and transcriptome assembly with assistance from KAF. ST characterised the AMPs with assistance from KAF and EP. ST drafted the manuscript with assistance from KAF. All authors provided review and feedback.

