## Supplementary material for "Multi-omics resources for the Australian southern stuttering frog (*Mixophyes australis*) reveal assorted antimicrobial peptides"

Table S1. Genomic elements in the southern stuttering frog (*M. australis*) genome. Reported in this table are; short interspersed nuclear elements (SINEs), divided into mammalian-wide interspersed repeats (MIRs); long interspersed nuclear elements (LINEs), divided into LINE1, LINE2 and L3/CR1 elements; long terminal repeat (LTR) elements, divided into endogenous retrovirus (ERVs) families ERV_classI, ERV_classII, and endogenous retrovirus-like (ERVLs); DNA elements, divided into DNA transposons hAT-Charlie and TcMar-Tigger; unclassified repeat elements; as well as small RNA and satellite repeats.

| **Genomic Element** | **Number of Elements** | **Length Occupied (bp)** | **% of genome** |
| --- | --- | --- | --- |
| SINEs: | 3679 | 723762 | 0.02 |
| MIRs | 1638 | 171279 | 0.01 |
| LINEs: | 469153 | 218952392 | 7.00 |
| LINE1 | 90324 | 57623295 | 1.84 |
| LINE2 | 265513 | 108665523 | 3.47 |
| L3/CR1 | 61919 | 23943509 | 0.77 |
| LTR Elements: | 595518 | 369662457 | 11.82 |
| ERVL | 717 | 389114 | 0.01 |
| ERV_classI | 53274 | 58971789 | 1.89 |
| ERV_classII | 770 | 147971 | 0.00 |
| DNA Elements: | 918500 | 326232044 | 10.43 |
| hAT-Charlie | 65103 | 18933154 | 0.61 |
| TcMar-Tigger | 201 | 109673 | 0.00 |
| Unclassified: | 3479824 | 764250609 | 24.43 |
| Total interspersed repeats: | N/A | 1679821264 | 53.70 |
| Small RNA: | 23893 | 21775567 | 0.70 |
| Satellites: | 8319 | 1441327 | 0.05 |

Table S2**.** Benchmarking Universal Single-Copy Orthologues (BUSCO) v5.3.2 scores from *M. australis*, *X. tropicalis, X. laevis, L. dumerilii*, and *P. corroboree* using the vertebrata_odb10 lineage*.* ‘C’ refers to the proportion of intact vertebrate orthologues identified in the nuclear genome, the transcriptome, or the annotated gene sets. These were then identified as either ‘S’=single copy or ‘D’=duplicated. ‘F’ refers to the proportion of fragmented orthologues discovered, and ‘M’ is the proportion of orthologues that were missing from the resource. There was no publicly available transcriptome for *L. dumerilii, P. ornatum* and *P. corroboree*, so transcriptome BUSCOs could not be generated for these species. There was no gene annotation available for *P. ornatum* and *P. corroboree*, so protein BUSCOs could not be generated for these species. The NCBI RefSeq or GenBank assembly number is provided.

| **Data** | **Species** | **BUSCO Scores** |
| --- | --- | --- |
| Genome | Southern stuttering frog (*M. australis*) | C:91.8%[S:90.3%,D:1.5%],F:2.9%,M:5.3% |
|  | African clawed frog (*X. laevis*) [GCF_017654675.1] | C:95.9%[S:60.5%,D:35.4%],F:1.1%,M:3.0% |
|  | Western clawed frog  (*X. tropicalis*) [GCF_000004195.4] | C:95.3%[S:94.5%,D:0.8%],F:1.6%,M:3.1% |
|  | Southern banjo frog (*L. dumerilii*) [GCA_011038615.1] | C:80.9%[S:79.9%,D:1.0%],F:9.8%,M:9.3% |
|  | Ornate burrowing frog  (*P. ornatum*) [GCA_016617825.1] | C:42.4%[S:41.8%,D:0.6%],F:25.4%,M:32.2% |
|  | Corroboree frog (*P. corroboree*) [GCA_028390025.1] | C:93.1%[S:87.9%,D:5.2%],F:3.2%,M:3.7% |
| Transcriptome | Southern stuttering frog  (*M. australis*); Gonads | C:93.0%[S:38.4%,D:54.6%],F:3.2%,M:3.8% |
|  | Southern stuttering frog  (*M. australis*); Liver | C:84.0%[S:50.4%,D:33.6%],F:5.6%,M:10.4% |
|  | Southern stuttering frog  (*M. australis*); Spleen | C:90.2%[S:48.8%,D:41.4%],F:3.4%,M:6.4% |
|  | Southern stuttering frog  (*M. australis*); Brain | C:89.9%[S:49.0%,D:40.9%],F:4.5%,M:5.6% |
|  | Southern stuttering frog  (*M. australis*); Dorsal Skin | C:89.8%[S:49.7%,D:40.1%],F:3.5%,M:6.7% |
|  | Southern stuttering frog  (*M. australis*); Ventral Skin | C:89.3%[S:51.3%,D:38.0%],F:3.6%,M:7.1% |
|  | Southern stuttering frog  (*M. australis*); Global | C:96.1%[S:24.6%,D:71.5%],F:1.7%,M:2.2% |
|  | African clawed frog (*X. laevis*); Follicle [GJQV00000000.1] | C:70.8%[S:54.5%,D:16.3%],F:12.9%,M:16.3% |
|  | Western clawed frog  (*X. tropicalis*); Gonads [GIVH00000000.1] | C:95.7%[S:30.8%,D:64.9%],F:0.9%,M:3.4% |
| Gene Annotation | Southern stuttering frog  (*M. australis*) | C:86.0%[S:84.0%,D:2.0%],F:8.5%,M:5.5% |
|  | African clawed frog (*X. laevis*) [GCF_017654675.1] | C:99.0%[S:28.9%,D:70.1%],F:0.4%,M:0.6% |
|  | Western clawed frog  (*X. tropicalis*) [GCF_000004195.4] | C:98.6%[S:50.2%,D:48.4%],F:0.6%,M:0.8% |
|  | Southern banjo frog (*L. dumerilii*) [GCA_011038615.1] | C:85.5%[S:82.6%,D:2.9%],F:8.0%,M:6.5% |

Table S3**.** Genomic coordinates of stuttering-barred frog cathelicidins and defensins.

| **AMP** | **Scaffold** | **Start** | **Stop** | **Strand** |
| --- | --- | --- | --- | --- |
| MA-CATH1 | 5 | 288567711 | 288571049 | + |
| MA-CATH2 | 5 | 288586976 | 288590222 | + |
| MA-CATH3 | 5 | 288665422 | 288671071 | - |
| MA-CATH4 | 5 | 288691522 | 288700039 | + |
| MA-CATH5 | 5 | 288709815 | 288719070 | + |
| MA-CATH6 | 5 | 288790820 | 288798573 | + |
| MA-CATH7 | 5 | 288691522 | 288850431 | + |
| MA-CATH8 | 5 | 288893048 | 288901423 | + |
| MA-CATH9 | 5 | 289036596 | 289044229 | + |
| MA-CATH10 | 5 | 289072868 | 289083938 | + |
| MA-CATH11 | 5 | 289087411 | 289110797 | + |
| MA-CATH12 | 5 | 291238657 | 291259111 | - |
| MA-BD1 | 2 | 159917 | 173293 | + |
| MA-BD2 | 2 | 1389356 | 1394702 | - |

Table S4**.** Physiochemical properties of predicted cathelicidin and β-defensin mature peptides, from the southern stuttering frog (*M. australis*). The top tBLASTn hit against the NCBI nucleotide collection (nr/nt)database according to percentage identity (%) is also shown.

| **AMP** | | **Molecular Weight (kDa)** | **Charge at pH 7** | **Hydrophobicity (%)** | **Top tBLASTn hit** | **% Identity** |
| --- | --- | --- | --- | --- | --- | --- |
| Cathelicidin | MA-CATH1 | 3833.20 | 8.9 | 17.07 | *Amolops loloensis* cathelicidin-AL mRNA, complete cds / JF923766.1 | 53.72 |
|  | MA-CATH2 | 2012.34 | 4.9 | 26.09 | *Amolops loloensis* cathelicidin-AL mRNA, complete cds / JF923766.1 | 52.59 |
|  | MA-CATH3 | 15170.17 | 27.3 | 12.42 | *Hyla sarda* RNA-binding protein cabeza-like (LOC130272483), mRNA / XM_056518320.1 | 54.84 |
|  | MA-CATH4 | 1905.14 | 0.9 | 36.84 | PREDICTED: *Rana temporaria* antimicrobial protein CAP18-like (LOC120941678), mRNA / XM_040355220.1 | 47.83 |
|  | MA-CATH5 | 2328.78 | 5.9 | 36.36 | *Odorrana livida* cathelicidin family antimicrobial peptide OL-CATH2 mRNA, complete cds / MH282907.1 | 43.24 |
|  | MA-CATH6 | 1344.53 | 0.9 | 53.85 | No homolog / N/A | N/A |
|  | MA-CATH7 | 1626.87 | 1.9 | 40 | *Hoplobatrachus rugulosus* cathelicidin mRNA, complete cds / MW725232.1 | 38.1 |
|  | MA-CATH8 | 1223.37 | -0.8 | 40 | No homolog / N/A | N/A |
|  | MA-CATH9 | 5927.17 | 2.9 | 43.14 | *Hyla sarda* cathelicidin-related antimicrobial peptide Bf-CRAMP-like (LOC130331777), partial mRNA / XM_056553719.1 | 43.33 |
|  | MA-CATH10 | 5198.03 | 4.1 | 39.58 | PREDICTED: *Rana temporaria* uncharacterized LOC120941693 (LOC120941693), mRNA / XM_040355248.1 | 39.13 |
|  | MA-CATH11 | 3708.29 | 1.1 | 34.38 | PREDICTED: *Bufo bufo* cathelicidin-4-like (LOC121000821), mRNA / XM_040431492.1 | 31.41 |
|  | MA-CATH12 | 5044.72 | 0.1 | 34.15 | PREDICTED: *Bufo bufo* retinoic acid receptor responder protein 2-like (LOC121000788), mRNA / XM_040431418.1 | 48.46 |
| Β-defensin | MA-BD1 | 5292.91 | 2.2 | 25.53 | PREDICTED: *Mauremys mutica* uncharacterized LOC123375363 (LOC123375363), ncRNA / XR_006581416.1 | 51.11 |
|  | MA-BD2 | 5219.94 | -2 | 42.55 | *Quasipaa spinosa* beta-defensin mRNA, complete cds / MZ889667.1 | 64.15 |

Table S5. List of publicly available frog cathelicidins and β-defensins as of the time of this study. These sequences were used in the homology-directed discovery of the southern stuttering frog AMPs, as well as the phylogenetic trees. The sequences for four fish cathelicidins used in figure 3 are also included.

| **AMP Name** | **Source Species** | **Whole Prepropeptide Sequence** | **Citation** |
| --- | --- | --- | --- |
| cathelicidin-AL | *Amolops loloensis* | MGLSATLWFLMGVAAGSMASPLLQWSEDDISVMALYSTDYYNKVSGEDVLYGLQENNTEYITDEKSRFHQLSFPIQKTVCQKSDNALTDDCAFKEGGVVKSCTSYFFEEDDRDIIVVTCQSQDGHREHSRVRRSRRGRGGGRRGGSGGRGGRGGGGRSGAGSSIAGVGSRGGGGGRHYA | (Hao et al. 2012) |
| cathelicidin-PY | *Paa yunnanensis* | MKVWQCVLWISALTLQMARSQSPDQDGWIREALDLYNQREDGEFFFKFLSDLPDVLLEEEGDSPAIGFLIKETDCPKSEDCDLEKCDYRKDGEVKVCTLYREEEDVKCVSLSKNSRTRRAGTKRKCNFLCKLKEKLRTVITSHIDKVLRPQG | (Wei et al. 2013) |
| Lf-CATH1 | *Limnonectes fragilis* | MKIWQCVLWLSAVTLARSQFLDQDGWIREALDLYNQREDGEYFFKVLSDVPADLLQEGEDSPEVAFLIKETECLKSEDNELARCDYKNDGEVKACGLYLEEGEASGTLKCVSLTKNPRMKRANRRPPCRGIFCRRVGSSSAIARPGKTLSTFITV | (Yu et al. 2013) |
| Lf-CATH2 | *Limnonectes fragilis* | MKIWQCVLWLSTVTWARSQSPDGDGWVREALDLYNQREDGEFYFKLLSDLPAALLQEEGDSPAVGFLIKETECRKSEAGDPEQCDYRQDGEVKVCALYREEEEVKCVSLTENSRIRRANKKGKCNVLCQLKQKLRSIGSGSHIGSVVLPRG | (Yu et al. 2013) |
| cathelicidin-RC1 | *Rana catesbeiana* | MKIWQCVVWLCAITLEVAHSQSPDREGWIREALDLYNQREDGEFLFKLLSELPGPLLEEEGDSPAIGFLIKETDCPKSEEIDLERCDYSKDGEVKVCALHQEEQDVKCVSLTENSRSKRSSKKKKCKFFCKVKKKIKSIGFQIPIVSIPFK | (Ling et al. 2014) |
| cathelicidin-RC2 | *Rana catesbeiana* | MKIWQCVVWLCAITLEVAHSQSPDREGWITEALDLYNQREDGEFLFKLLSELPAPLLEEEGDSPAIGFLIKETDCPKSEENDLEGCDYSKDGEVKVCALHREDEDVKCVSLTAENSRSKRSSNKKCGFFCKLKNKLKSTGSRSNIAAGTHGGTFRV | (Ling et al. 2014) |
| cathelicidin-PP1 | *Polypedates puerensis* | MKVWQCVIFLAALTLHLAHFQSGRIRAALDAYNQKDDGECYFKFVSDLPGSLLQEEEESPSVNFLIKETECRKSEDVDLERCEYKKDGEVKACALYLEKEEVECVSLSENSRPKRASENGKCNLLCLVKKKLRAVGNVIKTVVGKIA | (Mu et al. 2017) |
| BG-CATH37 | *Bufo gargarizans* | MRSWWLSLLLVSAVTLHGCLSDTAEPEVQDGRSVGDVIDLYNQREGVTYLYKSLDQLPPVPMEEDENPNRRGFIIKETVCLKSENPDLTQCDFKPDGDVKICSLDLDDEDPEDIMCTSLNKEVRVKRSSRRPCRGRSCGPRLRGGYTLIGRPVKNQNRPKYMWV | (Sun et al. 2015) |
| FM-CATH1 | *Fejervarya multistriata* | MKVWQCVLWICAITLHSARSQSSDQDGWIREALDLYNQKDDGEFCFKFLSDLPDALLEEEGDSQSIGFLIKETDCLKSEGQDLEQCDYKEDGEVKACVLSAEEEVKCVSLSEKRRTRRAIKKLKTKVLNKLKQKLQAVGNLIGSVIKA | (Wang et al. 2017) |
| FM-CATH2 | *Fejervarya multistriata* | MKVWQCVLWICAITLHSARSQSSDQDGWIREALDLYNQKDDGEFCFKFLSDLPDALLEEEGDSQSIGFLIKETDCLKSEGQDLEQCDYKEDGEVKACVLSAEEEVKCVSLSEKRRTRRAIKKLKTKALNKLKQKLQAVGNLIGSVIKG | (Wang et al. 2017; Wu et al. 2021) |
| Cathelicidin-OA1 | *Odorrana andersonii* | MGNYFQICVLFQAMTVLYSASIYHPLRRPLNSQLLEDSIKLFNEGLNNSSFLYKLLKVDSLKVDSHSQADSHENNSLTFVIKETVCPKAAEIPEECPFLEDGVMKNCTAMYSGPRQEKAELSCQTLDPPPQASSLPSTTSEGADEDQSSQSPHKAKFINSGKKNETAFLDVIGRDPTWSHLAASCLKCIFDDLPKTHN | (Cao et al. 2018) |
| cathelicidin-NV | *Nanorana ventripunctata* | MKVWQCALWISALTWQAARSQSPDREEWIREALDLYNQREDGEFFFKFLSDLPAAPLEEENNPTISFLIKETECLKSEDINLEECDYKKDGEVKVCGLYPEEGETMKTLKCVSLTKNFHTKRARGKKECKDDRCRLLMKRGSFSYV | (Wu et al. 2018) |
| camp | *Xenopus tropicalis* | MGATRMILVLASLAMAHIIWAFYLEDNRSVPLSSAFSRAIGLYNTQPGNKWAYKLLHIIPSSSRPRSTNPQIVRFTIKETSCARSVRRRLRKCPFKDGGLVKICSIAVSEDQDIVTVPVCQDVPEQRPSFASSEEESGSGEIIQDAKSRCRRPGSCTLIGRFNQRINRNQV | (Mitros et al. 2019) |
| OL-CATH1 | *Odorrana livida* | MEIWQCVIWLCAITLEVAHSQSPDQEEQIRAALGLYNQKEGGEFLFKLLSDLPAPLLQDGEDSPAVSFLIKETECLKSEDNNLAQCDYKKDGEVKICGLYTEEEEEETSEILKCVSLTENSRGERSKKKKKCKGYRCRPVGFSSPISRRINDSENIYLPFGV | (Qi et al. 2019) |
| OL-CATH2 | *Odorrana livida* | MEIWQCVIWLCAVTLEVAHSQSPVQEGRIREALDLYNQREDGEFLFKLLSELPASLLEEEGDSSAITFLIKETDCPKSEDNDLERCDYKEDGEVKVCALHREDEDVKCISLSENSRTKRSNKKRKCNFLCKVKNKLKSVGSKSLIGSATHHGIYRV | (Qi et al. 2019) |
| LOC108718684 | *Xenopus laevis* | MYPANMETTWGPILLLCLCLSVHGVSLPRIPMTDMEVSAMALVSTEYYNRGSSDNSIFQLFNSDIDHFTNGSSAQIQFTIKETGCQKSDNHMGQECDFLEDGVMKRCTASFFPEDELPAFVITCDAIQNPPLRVRRNGGRGRGGGRGGGIRGGGKSGFGSPIAGVKRNLINLPNAKIKRHFSA | (Fortriede et al. 2020) |
| LOC121394781 | *Xenopus laevis* | MAPGVISTHLILVCLAGMSSSAPLGVEKNDQLRSKIAKLIEFYNNQSSTDFLFKLLKVGDGTISDQQDGSTDAQTVSFVLQETDCFKLQEHTNPDNCVFKADGEVKQCSVSISAGEQELRCESPSALMHVQTKQWDQYSEALSKRLKSAECLECIFSQVPK | (Fortriede et al. 2020) |
| camp1.L | *Xenopus laevis* | MGVTWVILVLASLAMGQNIWAFYLEDNRSVPLFSPLSRAVDLYNTEQGNEWAFNLLDIIPSSSRPHSRNPRVVHFTIKETYCEGSVRKRLRKCSFKDGGLVKICSLTVFEDQNIVTVPVCDDVPEQLQQFNSESSKEESGSRQMIKDAKNRRKRPGSYSFIGRINRE | (Fortriede et al. 2020) |
| cathelicidin-DM | *Duttaphrynus melanostictu* | MRSWRLSLLLVSAVTLHGCLSDPAEPEVQDGRSIEDVIDLYNQREGVTYLYKSLDQLPPVPMEEDENPNRRGFIMKETVCLKSENPDLTQCDFKPDGDVKICSLDLGDEDPEDIMCFSLNKEVRMKRSSRRKPCKGWLCKLKLRGGYTLIGSATNLNRPTYVRA | (Shi et al. 2020) |
| PN-CATH1 | *Pelophylax nigromaculata* | MKVWQCVLWLCAITLDLAHSQPPDREDWIREALDLYNQREDGEFLFKLLSELPAPLLQEEGDSPAITFLIKETDCPKSEDNDLEGCDDKEDGEVKVCALHREEEDVKCVSLTENSRIKRSSNKKCNFFCKLKKKVKSVGSRNLIGSATHHHRIYRV | (Wang et al. 2021) |
| PN-CATH2 | *Pelophylax nigromaculata* | MKVWQCVLWLCAITLDLAHSQPPDREDWIRAALELCNQREDGEFLSKLLSELPTPPLQKEGDFPAIAFLIKETDCPKSKDNDLEGCDYKEDGEVKACAVQIEEEDVKCVSLSVGKMQKNSRIKRSSNEGCNILCLLKRKVKAVKNVVKNVVKSVVG | (Wang et al. 2021) |
| cath-MH | *Microhyla heymonsi* | MMGGCFWWGALLSAVTLQLSLCQTPGTGSSGEGLISEAVELYNQREDGEYLFQFLTQLPPILLEEEEDSPAVVFLIKETGCLKSEDKTSEECEYKEDGEVKICGLFPEENEEKIPLRCEDFSQESRPRRAAEGAPCKLGCKIKKVKQKIKQKLKAKVNAVKTVIGKISEHLG | (Chai et al. 2021) |
| HR-CATH | *Hoplobatrachus rugulosus* | MKVWQCAIWISAITLHLARSQSPDQDEWIREALDLYNQREDGEFFFKLLAGLPDDLLEDEGDSPSIGFLIKETDCPKVEDNDLEQCDYKKDGEVKVCALYGEEEEIKCYSLSKKSRTRRASKKGKCNLLCKLKQKLRSVGAGTHIGSVVLKG | (Chen et al. 2021) |
| LOC120941678_X2 | *Rana temporaria* | MKVWQCVVWLSAITLEVARSQSPDREGWIREAVDLYNQREDGEFLFKLLSELPAALLEEEGDSPAITFLIKETDCPKSEEKDLEGCDYKEDGEVKVCALQREEEDVKCVSLAEKSRSKRSSKKKCNFLCKVKNRLKSLSSTSVIAAGVPRGTYRG | (Streicher 2021) |
| PopuCATH | *Polypedates puerensis* | MALAAALTFLLGLACTILASPIQEWSEDDVAVMALYSADYYNKVSGEDVIYRLVGEEAEYIAGENVSFHQISFPIQETKCLKSDNKPTDDCAFKEGGVVKSCTSRFFEEDDRDVVVVTCQNQDGHQEHSRVRRSRGGRGGRGGGGSRGGRGSSGRGRTGSGSFIAGGGNRGSRGGRQYA | (Yang et al. 2022) |
| PopuDef | *Polypedates puerensis* | MKLFIMLLVFLGLVAITWGTSDGASPALWGCDSFLGYCRIACFAHEASVGQKDCAEGMICCLPNVF | (Wei et al. 2015) |
| defensin-TK | *Theloderma kwangsiensis* | LVFLVLVAITWGSSDGASPAIWGCDSFLGYCRLACFAHEASVGQKECAEGMLCCIPNVF | (Shen et al. 2016) |
| QS-BD | *Quasipaa spinosa* | MNSLTVLFLVLVAVTWTGGDGASPLIWGCENFLGYCRLACFARESSVGQKECAEGMLCCIPNVFGTFW | (Yu et al. 2022) |

| **AMP Name** | **Source Species** | **Whole Prepropeptide Sequence** | **Citation** |
| --- | --- | --- | --- |
| Cathelicidin 1 | *Orncorhynchus mykiss* | MKMKVQVRSLILLAVAVLQVRSQNQTETRYEDIILVALPQLLPGEEQAFRPILNQLQVKTLNTEDVDQSEVSVRLTFPIQETFCSKSQGQPGKPCPLKKNGKLMMCSMKVRHPILEASNNLNTDLSTFVCEYMDAEDALQQQIRTRRSKVRICSRDKNCVSRPGVGSIIGRPGGGSLIGRPGGGSVIGRPGGGSPPGGGSFNDEFIRDHSDGNRFA | (Chang et al. 2006) |
| Cathelicidin 2 | *Orncorhynchus mykiss* | MKMKALVRSLLLLAVANLLVRGQTQTETRYEDIITAASNQLLPVEEQAFHPLLNQLEVETLNTEDVDQSEVSVRLSFPLQETLCSKAQGQQGQPCPLKKNGKRMMCSMEVRHPILEIGNTLNTDRSDISCEYMEAEVQKIRTRRGKDSGGPKMGRKDSKGGWRGRPGSGSRLGFGSGIAGASGVNHVGTLPASNSTTHPLDNCKISPQ | (Chang et al. 2006) |
| Cathelicidin1 | *Salmo salar* | MKMKVQVRSLILLAVAVLQVRSQNQTETRYEDIILVALPQLLPGEEQAFRPILNQLQVKTLNTEDVDQSEVSVRLTFPMQETFCSKSPGQPGKPCPLKKNGKLMMCSMMVRHPILEASNNLNTDLSKTNCEYMEAEDAMQKIRTRRSQARKCSRGNGGKIGSIRCRGGGTRLGGGSLIGRLRVALLLGVAPFLLDLSQINVMEIAFA | (Chang et al. 2006) |
| Cathelicidin 2 | *Salmo salar* | MKMKALVRSLLLLAVASLLVRGQTQTETRYEDIITAASNQLLPVEEQAFHPLLNQLEVETLNTEDVDQSEVTVRLSFPLQETLCSKAQGQQGQPCPLKKNGKRMMCSMEVRHPILETGNTLNTDRSDISCEYMEAEDAMPVLSTQKIRTRRGKPSGGSRGSKMGSKDSKGGWRGRPGSGSRPGFGSSIAGASGRDQGGTRNA | (Chang et al. 2006) |

Table S6. Programs and packages used in the genome assembly, transcriptome assembly and genome annotation of the stuttering frog. Information on virtual machines (including processing power) and run times to the nearest 15 minutes are provided. When the same steps were run multiple times, the average runtime was recorded.

| **Analysis** | **Individual Steps** | **Virtual Machine Used** | **Program Used and Version** | **Run Time (HH:MM)** |
| --- | --- | --- | --- | --- |
| Genome Assembly | Filtering <Q20 DNA reads | Amazon Web Services r5.24x large machine (96 vCPU; 768 GB RAM) | bamtools v2.4.1 | <00:15 |
| Genome Assembly | Filtering adapter-containing reads | Amazon Web Services r5.24x large machine (96 vCPU; 768 GB RAM) | HiFiAdapterFilt v2.0.0 | 04:45 |
| Genome Assembly | Generating the unscaffolded, de novo genome assembly | Amazon Web Services r5.24x large machine (96 vCPU; 768 GB RAM) | hifiasm 0.19.2-r560 | 8:00 |
| Genome Assembly | Map Hi-C paired-end reads to assembled contigs | Pawsey Supercomputer (64 vCPU; 256GB RAM) | bwa-0.7.17-h7132678_9  picard-2.23.8-0  bedtools v2.30.0  samtools v1.15.1 | 12:00 |
| Genome Assembly | Merge mapped contigs into scaffolded, de novo genome assembly | The University of Sydney Artemis HPC (8CPUs and 50GB RAM requested) | YaHS v1.1  juicer_tools_1.19.02 | 59:30 |
| Genome Assembly | Mitochondrial genome assembly | The University of Sydney Artemis HPC (6CPUs and 20GB RAM requested) | MitoHiFi v3.2 | 1:00 |
| Genome Quality Assessment | Generating contig, scaffold and genome size statistics | The University of Sydney Artemis HPC (Terminal) | bbmap v38.86 | <00:15 |
| Genome Quality Assessment | Assessing gene completeness of genomes | Galaxy Australia Web Server | “Busco assess genome assembly and annotation completeness” (Galaxy Version 5.3.2+galaxy0) | 23:00 |
| Genome Quality Assessment | Generate k-mer databases from read files for merqury analysis | Pawsey Supercomputer (64 vCPU; 256GB RAM) | meryl v1.3 | 03:00 |
| Genome Quality Assessment | Assessing genome completeness and assembly accuracy using merqury | Pawsey Supercomputer (64 vCPU; 256GB RAM) | merqury v1.3  Java run time environment (JRE) v11.0.9.1  R v4.1.3  bedtools v2.30.0  samtools v1.15.1 | 01:00 |
| Genome Processing for Transcriptome Assembly | Generate database of repeat elements from genome | Pawsey Supercomputer (64 vCPU; 256GB RAM) | RepeatModeler v2.0.1 | 15:00 |
| Genome Processing for Transcriptome Assembly | Repeat mask the genome using the database of repeat elements | Pawsey Supercomputer (64 vCPU; 256GB RAM) | RepeatMasker v4.0.6 | 15:00 |
| Genome Processing for Transcriptome Assembly | Index the genome for faster RNA alignment | Pawsey Supercomputer (64 vCPU; 256GB RAM) | hisat2 v2.1.0 | 01:00 |
| Transcriptome Assembly | Trimming adapter sequences and low-quality RNA read segments | The University of Sydney Artemis HPC (4CPUs and 40GB RAM requested) | trimmomatic v0.39 | 26:00 |
| Transcriptome Assembly | Generating tissue-specific transcriptomes | The University of Sydney Artemis HPC (8CPUs and 80GB RAM requested) | hisat2 v2.1.0  samtools v1.9  stringtie v2.1.6 | 72:00 |
| Transcriptome Assembly | Merging tissue transcriptomes  into global transcriptome | The University of Sydney Artemis HPC (4CPUs and 40GB RAM requested) | TAMA v1.0  bedtools v2.29.2 | <00:15 |
| Transcriptome Assembly | Removal of low-expression and  non-coding transcripts | The University of Sydney Artemis HPC (4CPUs and 40GB RAM requested) | CPC2 v2019-11-19  bedtools v2.29.2 | 08:45 |
| Transcriptome Assembly | Prediction of coding regions from filtered global transcriptome | The University of Sydney Artemis HPC (1CPU and 20GB RAM requested) | TransDecoder v2.0.1  Perl v5.24 | 01:00 |
| Transcriptome Quality Assessment | Assessing gene completeness of transcriptomes | Galaxy Australia Web Server | “Busco assess genome assembly and annotation completeness” (Galaxy Version 5.3.2+galaxy0) | 01:00 |
| Genome Annotation | Gene prediction using FGENESH++ commercial software | Pawsey Supercomputer (64 vCPU; 256GB RAM) | FGENESH++ v7.2.2 | 26:45 |
| Genome Annotation Quality Assessment | Assessing gene completeness of transcriptomes | Galaxy Australia Web Server | “Busco assess genome assembly and annotation completeness” (Galaxy Version 5.3.2+galaxy0) | 00:30 |
| Genome Annotation Quality Assessment | Calculating gene statistics | The University of Sydney Artemis HPC (32CPUs and 5GB RAM requested) | genestats.sh  tabix v0.2.6  bedtools v2.29.2 | <00:15 |

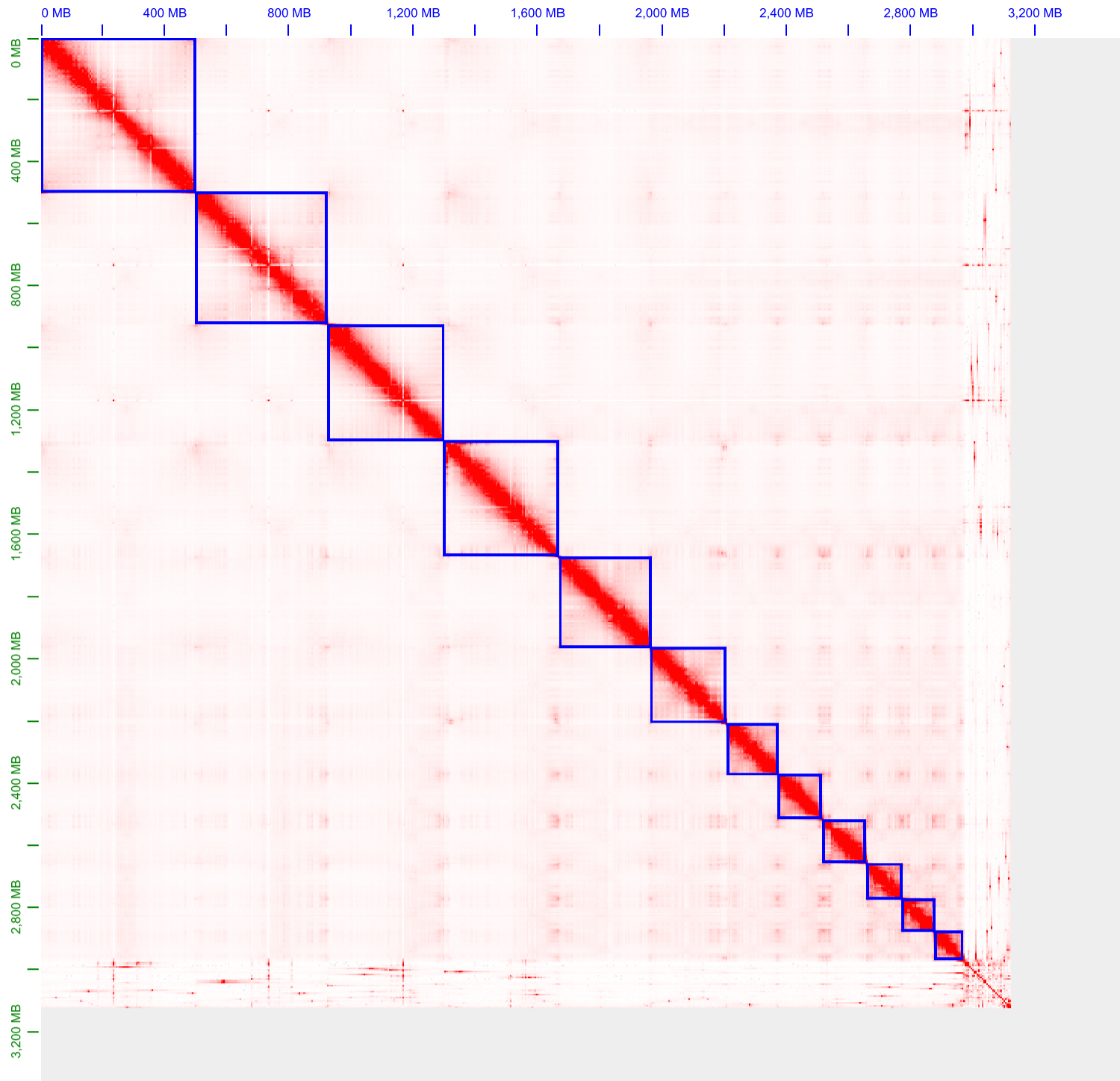

Figure S1. Hi-C contact map of the *M. australis* genome assembly. Red dots indicate interactions between regions of the genome, and blue boxes indicate scaffolds. Visualisation was generated using Juicebox v1.11.08 (Durand et al. 2016).

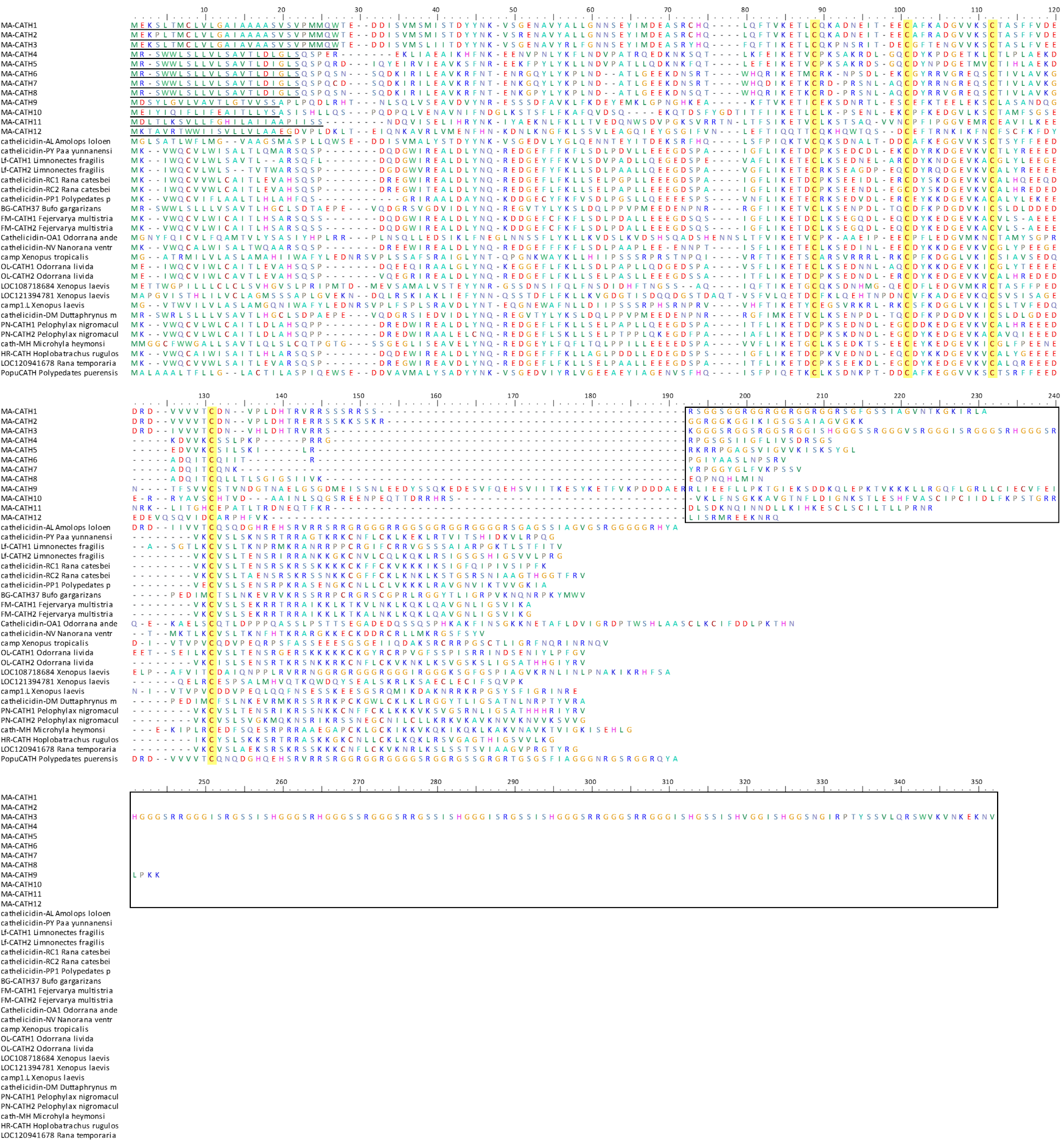
Figure S2. ClustalW multiple sequence alignment of putative stuttering frog cathelicidins (MA-CATH1 to 12) to known frog cathelicidins. For stuttering frog cathelicidins, the predicted signal peptide is underlined and predicted mature peptide boxed. The four highly conserved cysteine residues within the cathelin domain that are an identifying feature of this AMP family are highlighted in yellow.

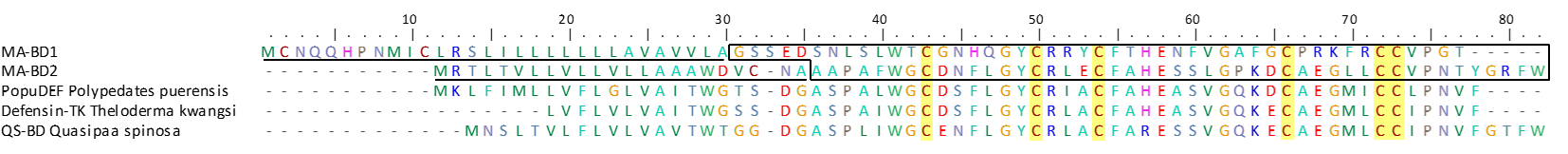
Figure S3. ClustalW multiple sequence alignment of stuttering frog β-defensins (MA-BD1 and MA-BD2) to other known frog β-defensins. For stuttering frog β-defensins, the predicted signal peptide is underlined and predicted mature peptide boxed. The six highly conserved cysteine residues within the mature peptide domain that are an identifying feature of this AMP family are highlighted in yellow.

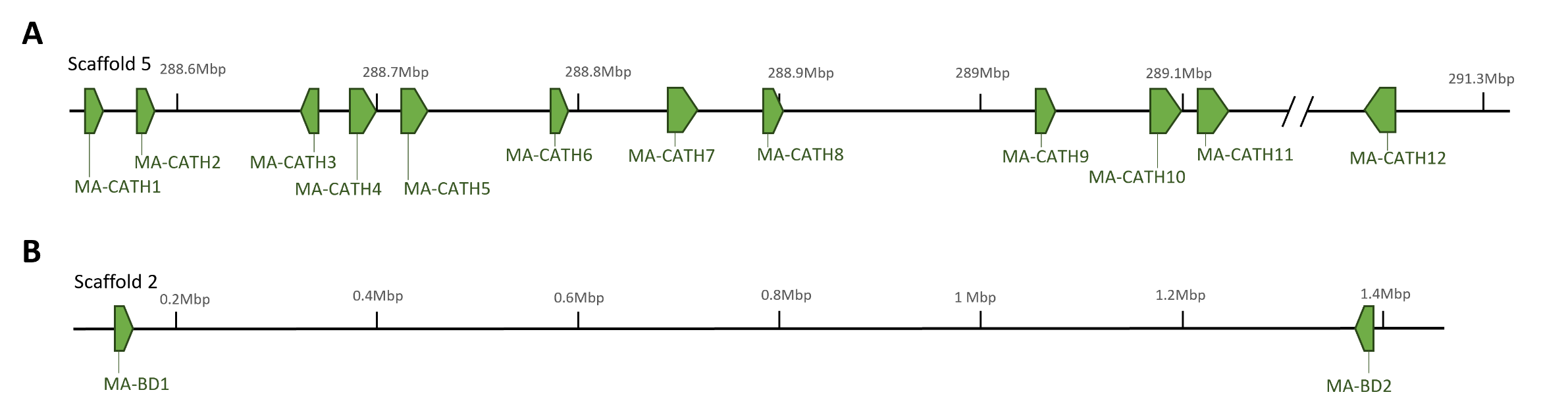

Figure S4**.** The genomic positions of the characterised southern stuttering frog **A)** cathelicidins on scaffold 5 and **B)** β-defensins on scaffold 2, with green arrows indicating the location of the gene and its direction.

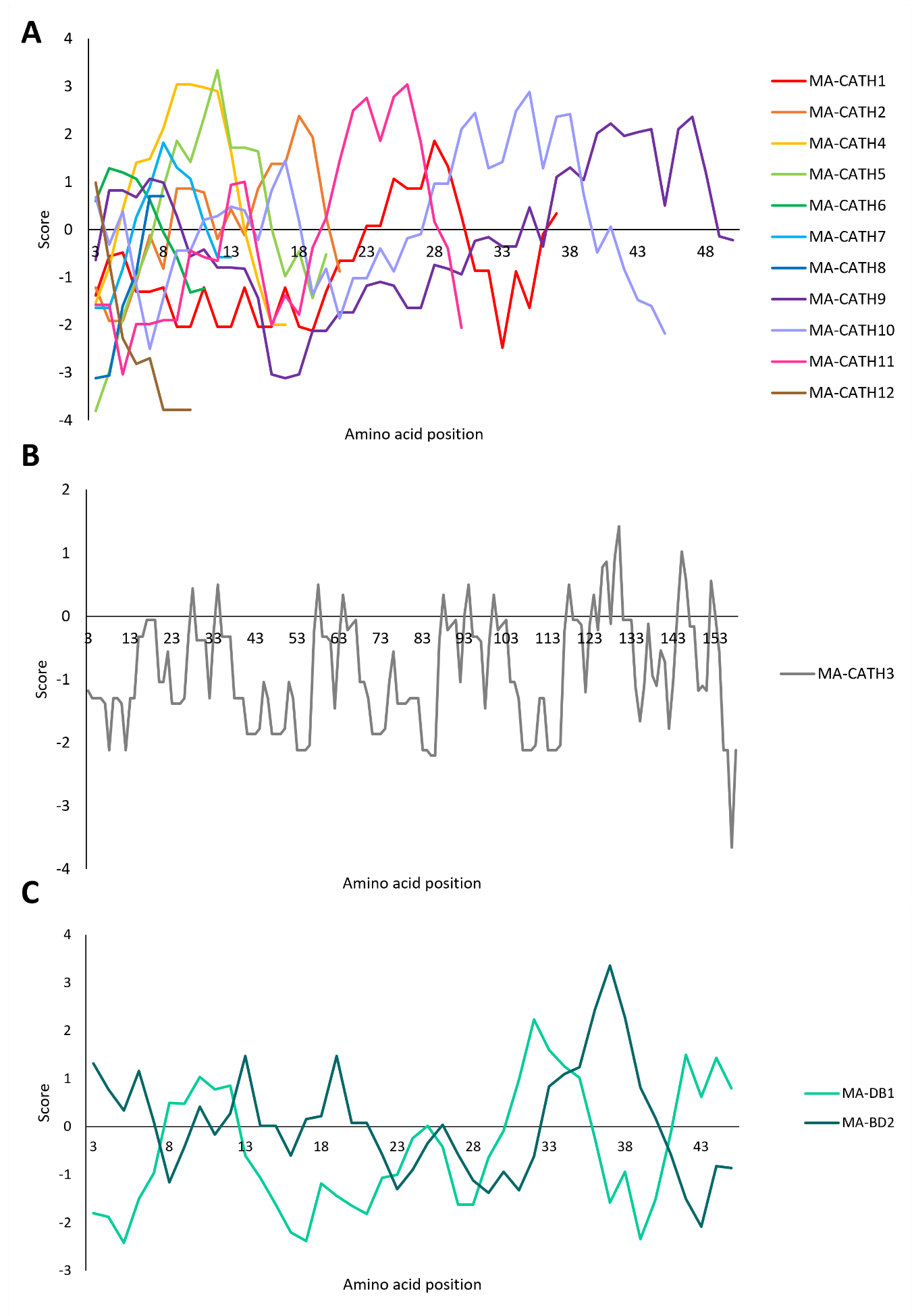
Figure S5**.** Kyte and Doolittle plots generated from the mature peptides of the southern stuttering frog cathelicidins (A-B) and β-defensins (C). Higher scores (>0) indicate hydrophobic regions, while lower scores (<0) indicate hydrophilic regions. Due to its length, MA-CATH3 was visualised separately.

### EXTENDED METHODS: Characterisation of stuttering frog cathelicidins and β-defensins

#### BLAST-directed AMP Discovery

To identify putative homologs of AMPs in the southern stuttering frog, we collated the protein sequences of all known frog cathelicidins and frog β-defensins from the DADP v1.6 (Novković et al. 2012) and the Antimicrobial Peptide Database v2022 (APD3) (Wang et al. 2016) using the search terms ‘cathelicidin’ and ‘defensin’. The National Center for Biotechnology Information (NCBI) Protein database was searched using all combinations of the query terms ‘frog’, ‘Anura’, ‘cathelicidin’, ‘CATH’, ‘defensin’, and ‘DEF’. Finally, Web of Science was searched using query terms ‘frog cathelicidin’ and ‘frog defensin’. The entire protein sequence was recorded for each AMP, and where available, the signal, pro-region, and mature sequences were also identified. All discovered frog cathelicidin and β-defensin sequences are in Table S5.

Using the entire prepropeptide sequences of these known frog AMPs as query sequences, a ‘tBLASTn’ search was used to search for homologs on the genome and global transcriptome. Any hits that have an e-value below a pre-defined threshold were then further evaluated. For cathelicidins, an e-value threshold of 0.01 was applied, while for β-defensins, (which have shorter, more variable sequences), the more lenient e-value of 10 was applied to avoid discarding possible matches. A BLASTp search was used to search for homologs against the annotated genes of the southern stuttering frog. BLASTp directly searches the protein query sequences against the protein sequences of the annotated genes. The same prepropeptide sequences and e-value thresholds were used.

#### HMMER-directed AMP Discovery

To characterise homologs of cathelicidins and β-defensins in the southern stuttering frog, multiple sequence alignments across a broad range of vertebrate species were downloaded from Pfam v35.0 (Mistry et al. 2021). For cathelicidins, a multiple sequence alignment of the highly conserved ‘cathelin’ region from various model species was downloaded from Pfam (ID: PF00666). Two multiple sequence alignments were available for the mature peptide of β-defensins, and they were both downloaded (ID: PF00711, PF13841).

To search against the global transcriptome, the iterative ‘jackhmmer’ function from HMMER was used. As HMMER can only iteratively search through protein sequence databases, the global transcriptome was first six-frame translated using EMBOSS v6.6.0.0 (Madeira et al. 2022). From there, jackhmmer was used to search the translated transcriptome against the Pfam regions, using the same e-value thresholds as the BLAST searches. Jackhmmer was also used to search the genome annotation.

#### Validating the Candidate AMP Genes

To confirm that the candidate transcripts and genes contained true cathelicidins and β-defensins and were not non-specific matches, all sequences passing the e-value thresholds were manually inspected for the presence of protein sequence motifs, as well as their genic structure.

Cathelicidins contain four, highly conserved cysteines, which in frogs, typically follow the spacing pattern **C**-X_9-11_-**C**-X_10_-**C**-X_9-16_-**C** (C=cysteine, X=any amino acid residue) (Chen et al. 2021). β-defensins generally have six, highly conserved cysteine residues, which in frogs, typically follow the spacing pattern **C**-X_6-9_-**C**-X_3-4_-**C**-X_9-13_-**C**-X_4-7_-**C-C** (Huttner and Bevins 1999; Machado and Ottolini 2015; Semple and Dorin 2012; Tang and Selsted 1993). To determine the presence of these sequence motifs in the putative AMP-encoding genes, the protein sequences of the candidate AMPs were aligned using the ClustalW algorithm against all the known frog AMPs of the corresponding peptide family (Figure S2; Table S5) and visually inspected. For candidate genes identified from the annotated genes, the protein sequence was already known, so they were directly aligned. However, for the matches solely identified from the genome or transcriptome, nucleotide sequences were first translated using ExPASy translate tool (Gasteiger et al. 2003), and the longest open reading frame was recorded. Any transcripts or annotated proteins that did not contain these structural motifs were removed from further analysis.

Cathelicidins are encoded by four exons, with the first three encoding the signal peptide and cathelin domain, while the fourth contains the proteolytic processing site, as well as the mature peptide (Tomasinsig and Zanetti 2005). β-defensins are expressed as either two-exon (Semple and Dorin 2012) or three-exon genes (Zou et al. 2007), with the first exon generally encoding the signal peptide, and the remaining exons encoding the mature peptide. To determine the number of exons of each candidate gene, several approaches were used. For candidate genes identified in the annotated genome, the number of exons is automatically calculated by FGENESH++ v7.2.2 (Softberry; Salamov & Solovyev, 2000; Solovyev et al., 2006; Tao et al., 2002). Candidate matches from the transcriptome were aligned against the genome using BLASTn. The transcript should align to the exonic regions of the corresponding gene, and the number of exons was confirmed by checking that the entire transcript is represented on the genome and that the exons appeared in sequential order, either on the forward or reverse strand. Any genes that did not have the expected number of exons were removed from further analysis.

The remaining candidate genes were manually annotated using Geneious Prime v2022.2 (https://www.geneious.com). The splice sites between each exon were inspected to determine if they exhibited traditional motifs, such as GT-AG, or GC-AG (Kitamura-Abe et al. 2004). Each candidate gene was inspected for premature stop codons. Any genes that did not exhibit the expected splice sites or stop codons were removed from further analysis. After these filtering processes, the remaining candidate genes were annotated as cathelicidin β-defensin genes.
